# Aerocavin is an antibiotic with potent and specific anti-Neisserial activity

**DOI:** 10.64898/2026.02.23.707490

**Authors:** Gleb Pishchany, Kyra E. Fryling, Vineetha Vasukuttan, Yern-Hyerk Shin, Tatum D Mortimer, Yonatan H. Grad, Jon Clardy

## Abstract

Gonorrhea, caused by *N. gonorrhoeae*, is a widespread sexually transmitted disease that is becoming resistant to all currently used antibiotics. Therefore, new therapeutics against gonorrhea are desperately needed. Here, we show that a natural product - aerocavin, is highly potent and specific against *Neisseria*. Aerocavin accumulates in *N. gonorrhoeae* at high levels and inhibits bacterial RNA polymerase (RNAP) by binding the switch region. Aerocavin resistance mutations evolve in *N. gonorrhoeae* at a low rate and are absent in clinical isolates. Previously overlooked narrow-spectrum antimicrobials like aerocavin may enable microbiome-sparing treatments of gonorrhea.

## INTRODUCTION

*Neisseria gonorrhoeae* is an obligate human pathogen that causes gonorrhea. Complications of the disease include pelvic inflammatory disease, ectopic pregnancy, female infertility, bacteremia, and an increased risk for HIV acquisition^1, 2^. More than 80 million new cases of gonorrhea are reported worldwide and over 600,000 in the United States ^3, 4^. Currently, the only recommended first-line antibiotic for the treatment of gonorrhea in the US is ceftriaxone^5^. Other countries also use azithromycin, cefixime, and ciprofloxacin, but resistance to all first-line antibiotics has been reported worldwide ^6–8^. High incidence and resistance require development of novel therapeutics against gonorrhea. Recently, two first-in-class antibiotics – zoliflodacin and gepotidacin, both targeting DNA gyrase – have shown non-inferiority to azithromycin/ceftriaxone combination therapy and approved by the FDA for treatment of uncomplicated urogenital gonorrhea ^9, 10^. No other first-in-class molecules that treat the disease are currently in the clinical pipeline.

Because *N. gonorrhoeae* is rarely included in primary antimicrobial screens, gonorrhea has been treated by broad-spectrum antibiotics developed for other infections. Recent reports have identified natural products with potent narrow-spectrum antimicrobial activity specifically against *Neisseria*^11–13^. These findings encourage revisiting antimicrobial molecules that may have been overlooked as candidates against *Neisseria* due to a narrow-spectrum of activity. Here, we report that aerocavin – a natural-product small molecule with modest activity against most pathogens - is highly active against *Neisseria* ^14^. We also identify aerocavin’s mechanism of action, quantify spontaneous resistance rates, and demonstrate increased accumulation in *N. gonorrhoeae* cells.

## RESULTS AND DISCUSSION

Aerocavin is a 14-carbon-ring macrolide with a decanoic acid side chain (Fig. 1A) produced by *Chromobacterium sp.* ATCC 53434^14^. A structurally similar antibiotic, ripostatin, with a phenylalkyl side chain instead of decanoate, is produced by *Sorangium cellulosum* (Fig. 1A)^15^. The biosynthetic gene cluster (BGC) for ripostatin, but not aerocavin, has been established^16^. Based on sequence homology to the ripostatin BGC, we have identified a candidate aerocavin BGC in the *Chromobacterium sp.* genome (Fig. 1B)^16^. Disrupting this putative BGC abolished aerocavin production (Fig. 1C), confirming its function.

**Figure 1.**
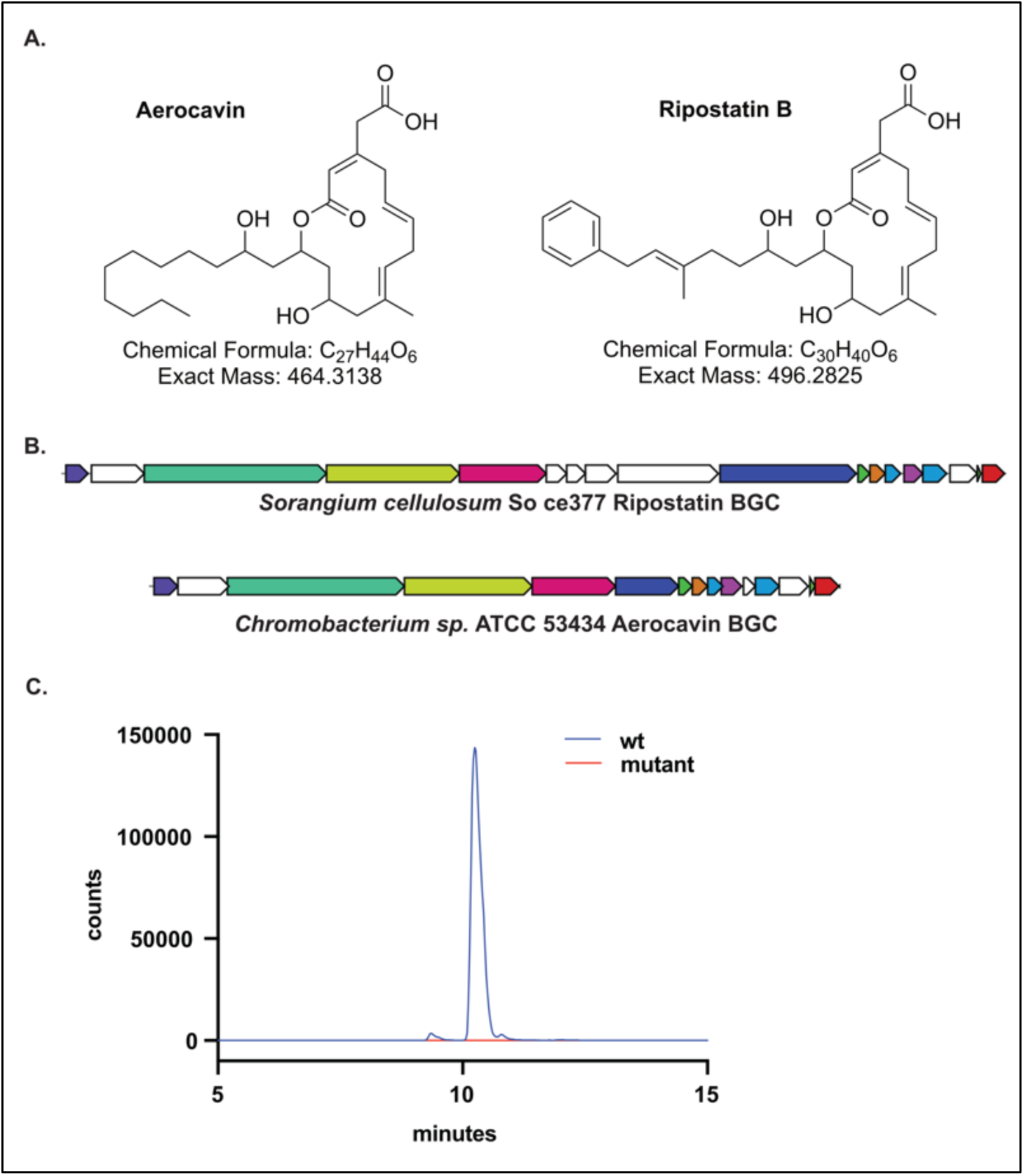
Aerocavin BGC is identified by sequence homology to ripostatin BGC. (A) Molecular structures of aerocavin and ripostatin. (B) Ripostatin BGC encoded by *S. cellulosum* So ce377 and predicted aerocavin BGC encoded by *Chromobacterium sp.* ATCC 53434. (C) Disruption of the putative aerocavin BGC eliminates aerocavin production by *Chromobacterium sp.* ATCC 53434.

We next assessed aerocavin’s activity against a panel of bacteria by disk-diffusion assays (Fig 2A). Aerocavin inhibited growth of methicillin-susceptible and methicillin-resistant *Staphylococcus aureus* (MSSA and MRSA), *N. gonorrhoeae*, and *Acinetobacter baumannii* but did not target other species (Fig. 2A). While aerocavin did not inhibit the wildtype *E. coli*, it prevented the growth of an isogenic *tolC* knockout strain. TolC is a component of AcrAB/TolC outer membrane efflux protein complex that pumps out diverse small molecules from the cytoplasm of enterobacteria^17^. This result suggests that *E. coli* resists aerocavin by active efflux.

**Figure 2.**
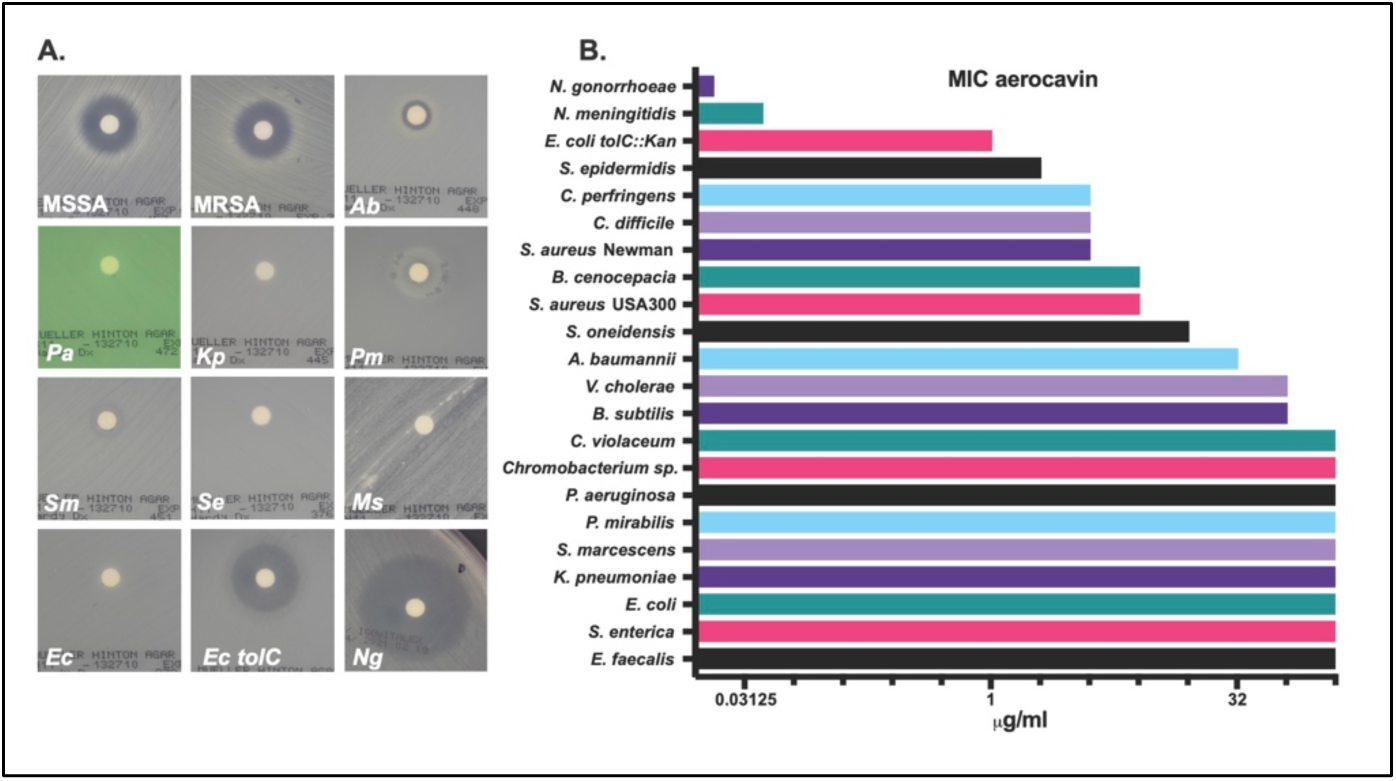
Aerocavin selectively inhibits *Neisseria*. (A) Disk-diffusion assay. MSSA – methicillin-susceptible *Staphylococcus aureus*, MRSA – methicillin-resistant *S. aureus*, *Ab* – *Acinetobacter baumannii*, *Pa – Pseudomonas aeruginosa, Kp – Klebsiella pneumoniae, Pm – Proteus mirabilis, Sm – Serratia marcescens, Se – Salmonella enterica, Ms – Mycobacterium smegmatis, Ec – Escherichia coli, Ng – Neisseria gonorrhoeae.* (B) Aerocavin’s minimal inhibitory concentrations.

We quantified aerocavin’s minimal inhibitory concentrations (MICs) against an expanded panel of bacteria. The MICs for most organisms ranged between 2 and >64 μg/mL (the highest concentration tested) (Fig 2B). Consistent with the disk diffusion assays, aerocavin inhibited the *tolC* knockout *E. coli* at 1 μg/mL, while the wildtype strain proliferated at 64 μg/mL. We noted that *N. gonorrhoeae* and *N. meningitidis* were inhibited at 0.02 and 0.04 μg/mL of aerocavin respectively – concentrations that are at least 100-fold lower than those required to inhibit other tested bacteria. Although ripostatin’s potency against *Neisseria* has not been reported, its activity against other tested microorganisms is in the same range as aerocavin’s^15, 18^. To define aerocavin’s spectrum of activity against clinically relevant *N. gonorrhoeae* strains we measured the MICs of aerocavin against a diverse panel of isolates (Table 1). None of the tested isolates, including those resistant to first-line antibiotics, had aerocavin MICs above 0.2 μg/mL.

**Table 1.** Minimal inhibitory concentrations of *N. gonorrhoeae* isolates.

| Isolate | ACV MIC<br>(µg/mL) | CRO MIC<br>(µg/mL) | CIP MIC<br>(µg/mL) | AZM MIC<br>(µg/mL) |
| --- | --- | --- | --- | --- |
| FA1090 | 0.025 |  |  | 0.25 |
| FA19 | 0.025 |  |  | 0.125 |
| 28B1 | 0.05 |  |  | 0.094 |
| FC428 | 0.1 | 0.5<sup>19</sup> | >32<sup>19</sup> | 0.25<sup>19</sup> |
| GCGS0457<br>*rpoD1* | 0.05 | 0.125<sup>20</sup> | 0.015<sup>20</sup> | 0.25<sup>20</sup> |
| GCGS0457<br>*rpoD2* | 0.05 | 0.19<sup>20</sup> | 0.015<sup>20</sup> | 0.25<sup>20</sup> |
| GCGS0745 | 0.2 | 0.03<sup>21</sup> | 0.015<sup>21</sup> | 256<sup>21</sup> |
| GCGS0394 | 0.025 | 0.008<sup>21</sup> | 0.015<sup>21</sup> | 2<sup>21</sup> |
| GCGS0604 | 0.1 | 0.032<sup>21</sup> | 0.016<sup>21</sup> | 16<sup>21</sup> |
| GCGS0473 | 0.05 | 0.008<sup>21</sup> | 0.015<sup>21</sup> | 2<sup>21</sup> |
| GCGS0138 | 0.2 | 0.008<sup>21</sup> | 0.015<sup>21</sup> | 16<sup>21</sup> |
| GCGS0807 | 0.1 | 0.06<sup>21</sup> | 16<sup>21</sup> | 0.5<sup>21</sup> |
| NZ2015-388 | 0.1 | 0.004<sup>22</sup> | 0.004<sup>22</sup> | 1<sup>22</sup> |
| GCGS0857 | 0.1 | 0.015<sup>21</sup> | 0.015<sup>21</sup> | 0.5<sup>21</sup> |
| GCGS0698 | 0.2 | 0.008<sup>21</sup> | 0.015<sup>21</sup> | 2<sup>21</sup> |
| GCGS0576 | 0.1 | 0.015<sup>21</sup> | 0.008<sup>21</sup> | 1<sup>21</sup> |
| GCGS0759 | 0.1 | 0.063<sup>21</sup> | 16<sup>21</sup> | 8<sup>21</sup> |
| GCGS0439 | 0.1 | 0.008<sup>21</sup> | 0.004<sup>21</sup> | 0.5<sup>21</sup> |
| GCGS0465 | 0.0125 | 0.008<sup>21</sup> | 0.015<sup>21</sup> | 0.06<sup>21</sup> |
| GCGS0781 | 0.0125 | 0.016<sup>21</sup> | 4<sup>21</sup> | 0.06<sup>21</sup> |
| GCGS0348 | 0.025 | 0.008<sup>21</sup> | 0.015<sup>21</sup> | 0.06<sup>21</sup> |
| NY0107 | 0.0125 | 0.002<sup>23</sup> | 0.002<sup>23</sup> | 0.016<sup>23</sup> |
| NY0609 | 0.0125 | 0.016<sup>23</sup> | 1.5<sup>23</sup> | 0.032<sup>23</sup> |

Abbreviations: ACV – aerocavin, CRO – ceftriaxone, CIP – ciprofloxacin, AZM – azithromycin.

*S. aureus* and *N. gonorrhoeae* are the most susceptible to aerocavin Gram-positive and Gram-negative species, respectively. To establish the ability of these pathogens to evolve spontaneous resistance to aerocavin, we measured the frequency of emergence of resistant mutants at a range of aerocavin concentrations. In *S. aureus*, the frequency of resistance was high and comparable to that of rifampicin (Fig 3A), but *N. gonorrhoeae* evolved resistance to aerocavin at lower rates (Fig. 3B). No resistant *N. gonorrhoeae* mutants were observed at concentrations above 4-fold of the wildtype aerocavin MIC (Fig 3B). These results indicate that in addition to being highly susceptible, *N. gonorrhoeae* has limited capacity to evolve resistance to aerocavin.

**Figure 3.**
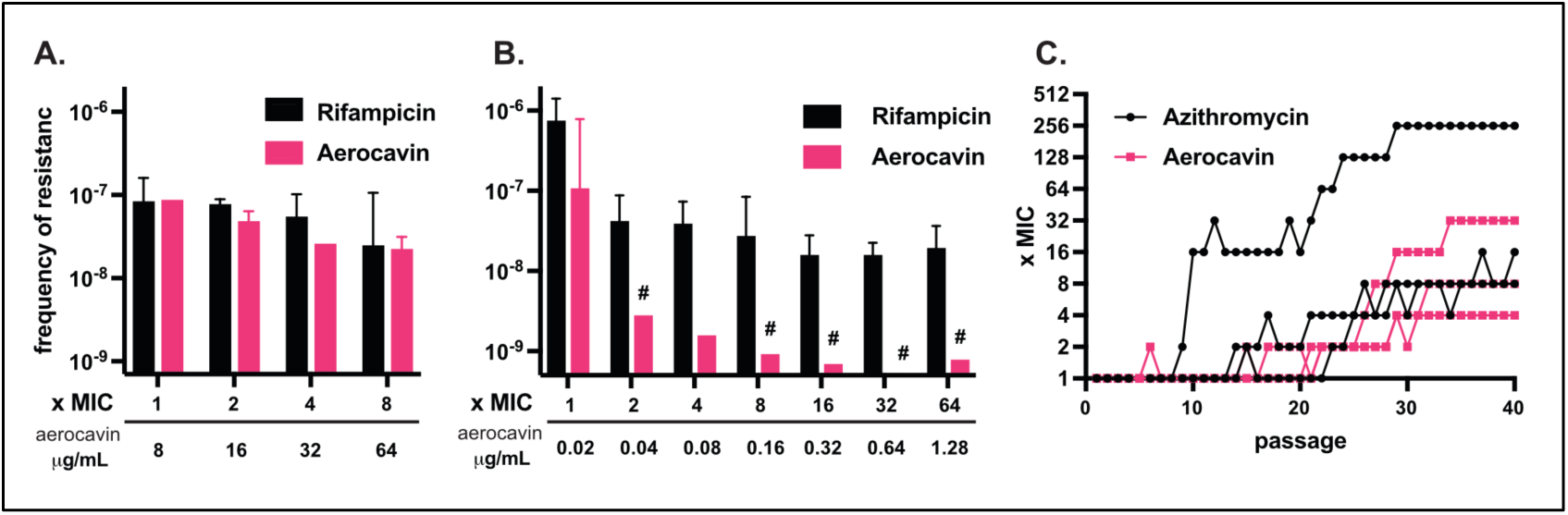
Resistance to aerocavin evolves in *N. gonorrhoeae* at low rates. Frequency of resistant mutants to aerocavin and rifampicin in (A) *S. aureus* and (B) *N. gonorrhoeae* at a range of concentrations on agar. No aerocavin-resistant mutants were recovered under conditions marked with “#” where the columns represent the limit of detection. (C) MICs of aerocavin and azithromycin in 3 replicates of *N. gonorrhoeae* evolved in broth over 40 passages.

To gain insight into aerocavin’s mechanism of action, we isolated aerocavin-resistant mutants of *S. aureus* and *N. gonorrhoeae* and sequenced their genomes. All resistant *S. aureus* isolates contained mutations in *rpoB* (encoding RNA polymerase subunit β) or *rpoC* (encoding RNA polymerase subunit β′) (Table S1) and were resistant to aerocavin at 64 μg/mL - at least a 16-fold increase in resistance over wildtype. Similarly, aerocavin-resistant *N. gonorrhoeae* with high levels of resistance (> 4-fold increase in the MIC) contained mutations in *rpoB* or *rpoC* (Table 2). Mutations conferring low-level resistance to aerocavin in *N. gonorrhoeae* localized to *mtrR*, encoding a repressor of the MtrCDE pump associated with multidrug resistance, *lptA*, encoding a putative LPS transport periplasmic protein altering outer membrane permeability^24^, and an uncharacterized putative acyl-CoA/acyl-ACP dehydrogenase (NGO_RS03140).

**Table 2.**
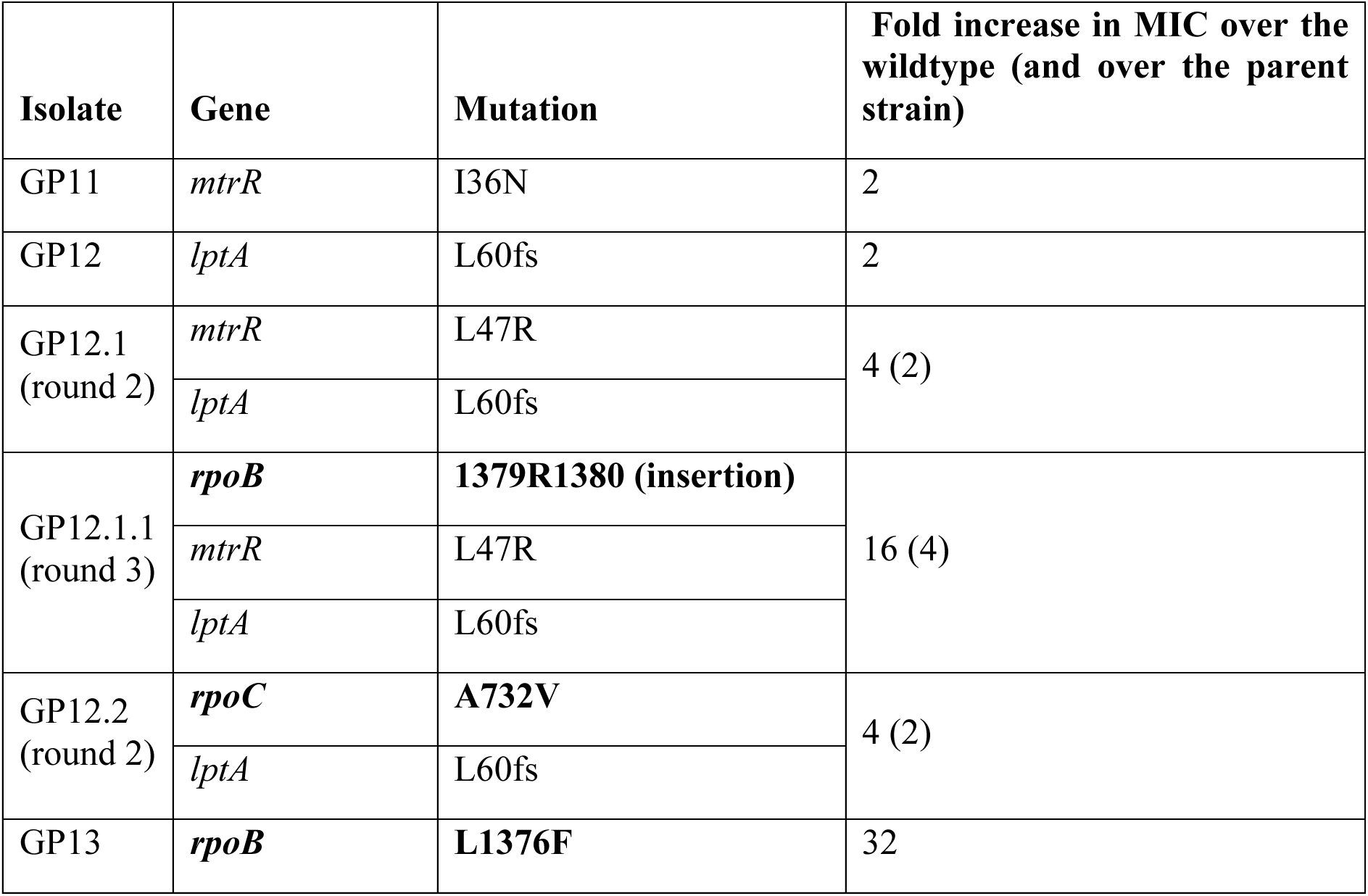
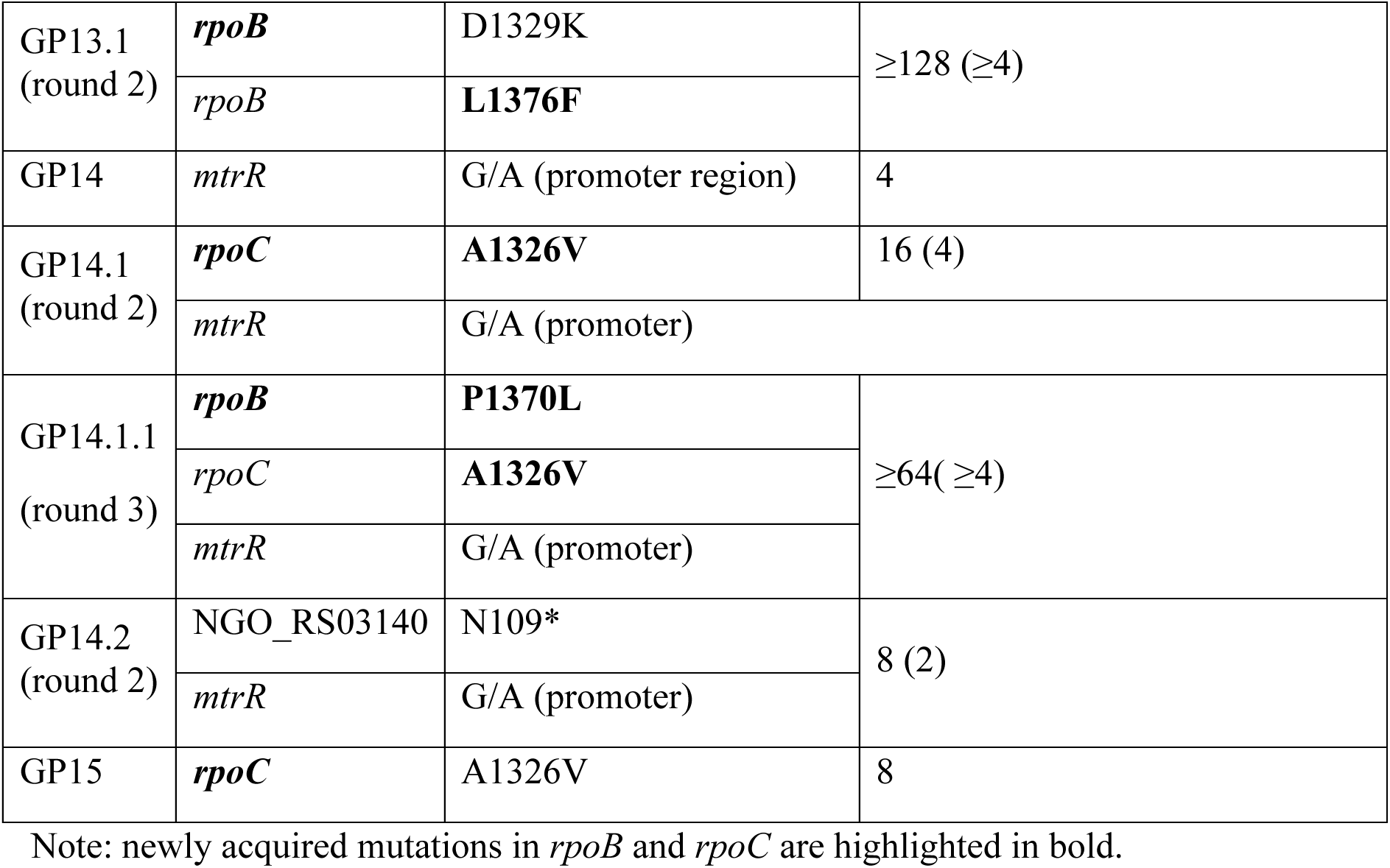
Aerocavin-resistant *N. gonorrhoeae* selected on agar.

| Isolate | Gene | Mutation | Fold increase in MIC over the wildtype (and over the parent strain) |
| --- | --- | --- | --- |
| GP11 | *mtrR* | I36N | 2 |
| GP12 | *lptA* | L60fs | 2 |
| GP12.1<br>(round 2) | *mtrR* | L47R | 4 (2) |
|  | *lptA* | L60fs |  |
| GP12.1.1<br>(round 3) | ***rpoB*** | **1379R1380 (insertion)** | 16 (4) |
|  | *mtrR* | L47R |  |
|  | *lptA* | L60fs |  |
| GP12.2<br>(round 2) | ***rpoC*** | **A732V** | 4 (2) |
|  | *lptA* | L60fs |  |
| GP13 | ***rpoB*** | **L1376F** | 32 |
| GP13.1<br>(round 2) | ***rpoB*** | D1329K | ≥128 (≥4) |
|  | *rpoB* | **L1376F** |  |
| GP14 | *mtrR* | G/A (promoter region) | 4 |
| GP14.1<br>(round 2) | ***rpoC*** | **A1326V** | 16 (4) |
|  | *mtrR* | G/A (promoter) |  |
| GP14.1.1<br>(round 3) | ***rpoB*** | **P1370L** | ≥64 (≥4) |
|  | *rpoC* | **A1326V** |  |
|  | *mtrR* | G/A (promoter) |  |
| GP14.2<br>(round 2) | NGO_RS03140 | N109* | 8 (2) |
|  | *mtrR* | G/A (promoter) |  |
| GP15 | ***rpoC*** | A1326V | 8 |

Note: newly acquired mutations in *rpoB* and *rpoC* are highlighted in bold.

We subjected a subset of resistant *N. gonorrhoeae* mutants to two more rounds of aerocavin selection and re-isolated the mutants with further increases in resistance (round 2 and 3). Second-and third-round mutants contained additional mutations in the same set of genes indicating that the acquisition of several mutations can have a combined effect on resistance (Table 2). For example, isolate GP13.1 acquired a D1329K mutation in *rpoB* in addition to the L1376K mutation inherited from its parent GP13. The first mutation (L1376K) increased the MIC by 32-fold, while the second mutation (D1329K) further increased resistance by at least another 4-fold to a final ≥ 128-fold increase in the MIC over the wildtype. Similarly, GP14.1.1. in addition to mutations in *rpoC* and *mtrR* inherited from GP14.1 acquired a mutation in *rpoB* further increasing its resistance to aerocavin.

To explore gradual evolution of aerocavin resistance, we passaged three replicates of *N. gonorrhoeae* 40 times in the presence of either aerocavin or azithromycin in broth at sub-MICs (Fig 3C). We sequenced the final passages of the three aerocavin broth replicates as well as passage #28 from replicate 1. Similarly to agar, mutations in *rpoB*, *rpoC,* and *mtrR* were selected for by aerocavin in broth (Table S2). Additionally, we observed mutations in other genes, which could conceivably increase resistance to aerocavin. For example, all three cultures contained mutants in putative (p)ppGpp synthetase/hydrolase *spoT*. Changes in (p)ppGpp levels have been implicated in the emergence of antibiotic tolerance and modulation of RNA polymerase activity ^25, 26^. Two of the three cultures contained frameshift mutations in *pilQ,* which is involved in the biogenesis of type IV pili and serves as a channel for the translocation of antimicrobials^27^. One culture contained a mutation in *atpA* and another in *atpB,* which encode putative components of an ATPase complex. Reduced levels of ATP in the cells are linked to persistence^28, 29^.These data confirm that low-level resistance to aerocavin evolves through mutations in genes involved in general response to stress and antibiotics, while high-level resistance requires mutations in RNA polymerase. To search for these aerocavin-resistant RNAP mutants within clinical isolates we scanned the genome sequences of over 12,000 isolates’ *rpoB* and *rpoC* sequences. None of the available genomes contained mutations that would provide high-level resistance to aerocavin identified here (Tables 2, S1, and S2). These results suggest that high-level resistance to aerocavin in *N. gonorrhoeae* is currently not present in clinical isolates although it is possible that other unidentified resistant mutations were missed.

Mapping of the mutations conferring high-level resistance to *rpoB* and *rpoC* implicates bacterial RNAP as aerocavin’s target. This is consistent with RNAP being inhibited by ripostatin^30^. Accordingly, an *in vitro* assay demonstrated that aerocavin inhibits *E. coli* RNAP with an IC_50_ of 78 nM (55-110 nM, 95% CI) (Fig 4A). Mutations conferring resistance to both aerocavin and ripostatin map to the switch region of bacterial RNA polymerase (Tables 2, S1, and S2)^31^, which together with their structural similarity (Figure 1A) suggests that that they have the same mechanism of action ^31^. Notably, structurally distinct fidaxomicin targeting the switch region of RNA polymerase, is used as a narrow-spectrum antibiotic against *Clostridium difficile*, validating the RNAP switch region as a target for antibiotic development^32^.

**Figure 4.**
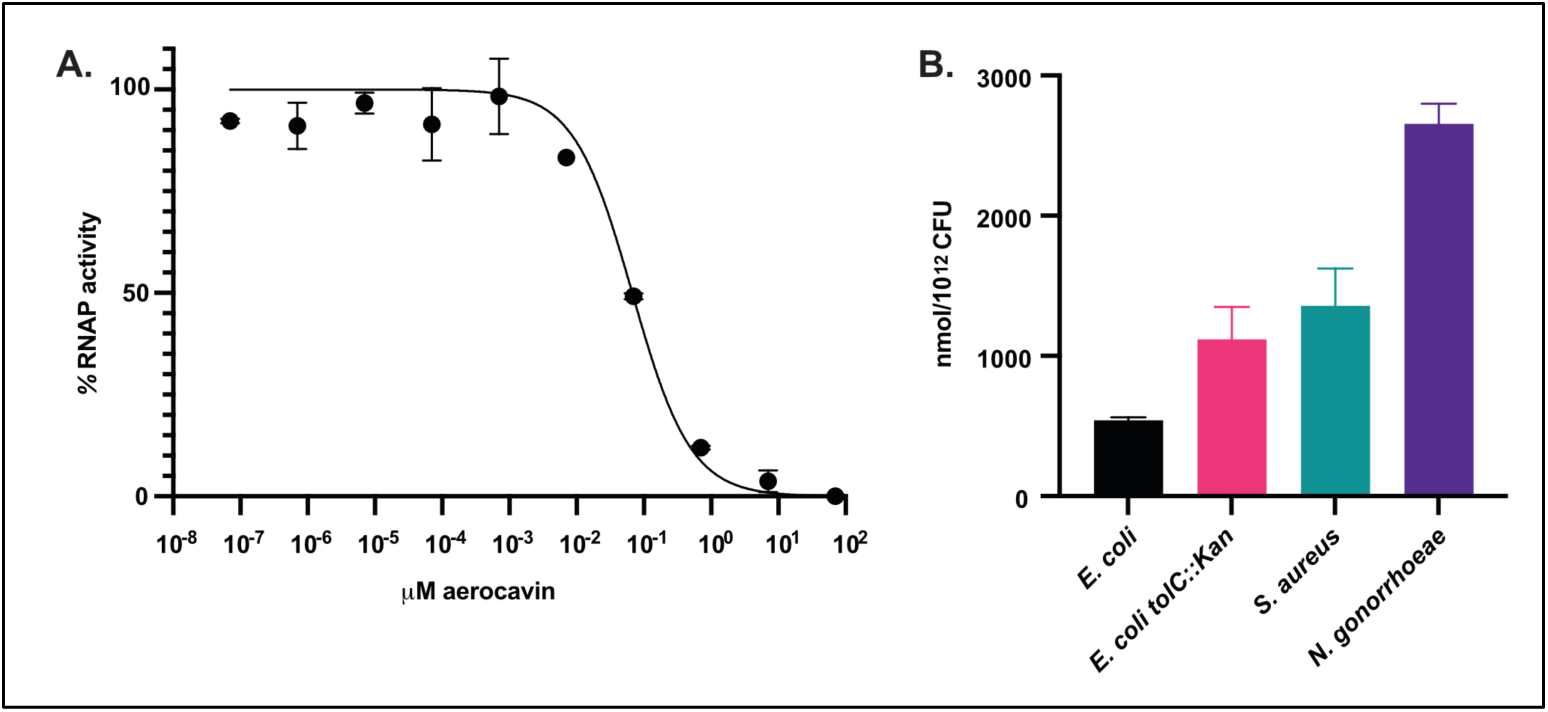
Aerocavin is a potent inhibitor of bacterial RNA polymerase with increased penetration of *N. gonorrhoeae* cells. (A) *In vitro* inhibition of *E. coli* RNAP by aerocavin. (B) Accumulation of aerocavin in bacterial cells.

We wondered whether aerocavin’s low MIC is due to its increased accumulation in *Neisseria* cells – a property reported for other small molecules^11, 12^. We measured the accumulation of aerocavin in wildtype and isogenic *tolC* knockout *E. coli*, *S. aureus*, and *N. gonorrhoeae* (Fig 4B).

*N. gonorrhoeae* accumulated aerocavin at higher concentrations than *E. coli* and *S. aureus*, and the *tolC* knockout *E. coli* accumulated more aerocavin than its wildtype counterpart (Figure 4B). These data are consistent with the susceptibilities of these strains to aerocavin (Fig. 2) and indicate that aerocavin’s specificity for *Neisseria* is at least partially due to its high intracellular accumulation.

Rising resistance and the dearth of new antibiotic classes have narrowed treatment options for gonorrhea. Most antimicrobials are identified and prioritized by screens in fast growing organisms (e.g., *E. coli* and *S. aureus*), while *N. gonorrhoeae* is usually omitted. This approach hampers discovery of agents specific to *N. gonorrhoeae* and other fastidious pathogens. Recent reports have identified natural products and synthetic molecules with narrow-spectrum activity against *N. gonorrhea* and other pathogens ^11–13, 33–35^. Interest in narrow-spectrum antimicrobials is driven both by challenges in developing novel antimicrobial agents and by recognition of collateral damage to microbiota caused by broad-spectrum antibiotics^36, 37^. A practical hurdle to clinical use of narrow-spectrum antibiotics is that therapy often starts before the identification of the causative agent. However, diagnosis of gonorrhea is straightforward and rapid, making it a good candidate for narrow-spectrum treatments. Innovative approaches to rapid point-of-care diagnostics may make the use of narrow-spectrum precision antibiotics a reality for other infections^38^.

## METHODS

### Bacterial strains and culturing conditions

All strains and growth conditions are listed in Table S3. *Chromobacterium sp.* ATCC 53434, *N. gonorrhoeae* FA1090, *S. aureus* Newman, and *E. coli* K-12 were used unless otherwise noted.

### Identification and validation of aerocavin BGC

Putative aerocavin BGC was identified by antiSMASH^39^. To disrupt the BGC, 1 kb regions flanking the first gene in the operon (CXB49_16090, homologue of RipA, predicted to encode a 3-hydroxyl-3-methylglutaryl-CoA synthase) were amplified using the acvA1_fwd/acvA1_rev and acvA2_fwd/acvA2_rev primer pairs listed in Table S4. The PCR products were joined and inserted into pNPTS138 Cm plasmid (amplified with pNPTS138_fwd/pNPTS138_rev primers) using NEBuilder HiFi DNA Assembly Cloning Kit (NEB). The construct was transformed into *E. coli* S17-1 and transformants were selected on LB agar supplemented with 20 μg/mL chloramphenicol. To carry out the conjugation, S17-1 was grown in LB broth supplemented with chloramphenicol and *Chromobacterium sp.* ATCC 53434 was grown in LB until early stationary phase. Bacterial cells were washed in LB and resuspended in 1/10^th^ volume of fresh LB broth. One hundred μL of S17-1 were mixed with 10 μL of ATCC 53434 and 20 μL of the mix were spotted onto LB agar and incubated at 30 °C overnight. Cells were then scraped, resuspended in 100 μL LB broth, and spread on LB agar containing 20 μg/mL chloramphenicol (to select for the plasmid) and 100 μg/mL ampicillin (to select for ATCC 53434) and incubated overnight at 30 °C. Because of frequent spontaneous resistance to chloramphenicol, multiple colonies were picked, incubated overnight with shaking in LB at 30 °C, and screened for the presence of pNPTS138 Cm plasmid (pNPTS_sacB_PCR1/ pNPTS_sacB_PCR2 primers, Table S4). The overnight cultures were diluted 10,000-fold and 100 μL were spread onto LB containing 5% (v/w) sucrose and incubated for 2 days. Multiple colonies were screened for the loss of chloramphenicol resistance without success. PCR indicated that the strains were not cured of the plasmid even after multiple passages on LB/sucrose. We reasoned that, even without double cross-over, plasmid integration within the BGC would disrupt its function. We set up a PCR to amplify the region downstream from the plasmid integration site using acvA-seq4/ M13_pUC_fwd primer pair (Table S4). acvA-seq4 is located within the chromosome downstream from acvA2 flanking region, while M13_pUC_fwd is located within pNPTS138 Cm. The resulting PCR product was sequenced and confirmed to contain the acvA1-acvA2 construct and to lack acvA gene. We additionally PCR-amplified the region upstream from the plasmid insertion site with acvA-seq1-1 primer (located within the chromosome) and M13_pUC_rev primer. Sequencing of the resulting PCR product confirmed that acvA flanked by acvA1 and acvA2 was upstream from the insertion site. Wild type and mutant strains were extracted with ethyl acetate and fractionated with C18 Sep-Pak columns (Waters) with stepwise acetonitrile gradient. The fraction eluted with 75% acetonitrile was analyzed by LCMS using an Agilent 1200 series HPLC system equipped with a photo-diode array detector and a 6130-quadrupole mass spectrometer on a Phenomenex Kinetex C18 100Å (100 x 4.6 mm) column under the following LC method: hold 50% ACN + 0.1 % FA/water + 0.1% FA for 1 minute then gradient to 100% ACN + 0.1% FA over 17 minutes then hold at 100% ACN + 0.1% FA for 1 minute with constant flow rate of 0.4 mL/minute.

### Determination of frequency of resistant mutants

One hundred microliters of an overnight culture of *S. aureus* were spread over tryptic soy agar (*S. aureus*) containing aerocavin or rifampicin. In parallel the culture was serially diluted and plated on tryptic soy agar without antibiotics for estimation of the total number of viable bacteria. The number of colonies growing at different concentrations of antibiotics were used to calculate the frequency of resistance. Frequency of resistance in *N. gonorrhoeae* was estimated similarly, except the cells were collected from chocolate agar and resuspended in Mueller-Hinton broth to an OD_600_ of ∼2.0 prior to plating on GCB agar (15g/L proteose peptone 3, 1 g/L soluble starch (Difco), 4g/L dibasic potassium phosphate, 1 g/L monobasic potassium sphosphate, 5g/L NaCl, 5g/L glucose) supplemented with 1% IsoVitaleX^TM^ solution.

### Evolution of resistance in liquid cultures

Three colonies of *N. gonorrhoeae* were inoculated into 1 mL of GCB broth supplemented with 1% IsoVitaleX^TM^ solution and grown overnight. Ninety-six well plates containing 150 μl of the same medium and supplemented with 2-fold dilutions of aerocavin or azithromycin (0.002 - 1 μg/mL) were inoculated from the overnights at 1:1,000 dilution. The plates were incubated for 24 hours at 37 °C in 5% CO_2_. After 24 hours, the plates were visually inspected for the presence of turbidity. The wells with the highest concentration of aerocavin or azithromycin that allowed growth were used to inoculate the next passage (1/100 dilution – 1.5 μl inoculum into 150 μl media). Two separate replicates were simultaneously passaged without the antibiotics serving as wildtype MIC controls. As the MICs of the passages increased, we elevated the concentrations of the supplemented antibiotics accordingly. After 20 passages the experiment was paused and later resumed with the following modifications aimed to accelerate evolution of resistance: the total volume was increased to 200 μl and the inoculum to 10 μl (1/20 dilution).

### Aerocavin uptake assay

Aerocavin uptake into bacterial cells was measured as previously described with slight modifications^40^. All species were grown in GCB broth at 37 °C without shaking at 5% atmospheric CO_2_ until they reached stationary phase (overnight for *S. aureus* and *E. coli*, two days for *N. gonorrhoeae*). These cultures were then diluted into 100 mL fresh media. Because *N. gonorrhoeae* did not proliferate in 100 mL volume, the culture was separated into 5 mL aliquots, which were combined immediately after growth. The cultures were allowed to reach optical density at 600 nm (OD_600_) of 0.4-0.6 (requiring overnight growth for *N. gonorrhoeae*), pelleted through centrifugation at 3,000 *g* for 10 minutes at 4 °C, washed in ice-cold phosphate buffered saline (PBS), pelleted again and resuspended in PBS to OD_600_ of 10. The suspensions were incubated with shaking at 37 °C for 5 minutes, and small aliquots were taken for measurement of CFU/mL through serial dilution and plating on chocolate agar. Aerocavin was added to 22 μM (10 μg/mL) and the suspensions were incubated at 37 °C with shaking for another 10 minutes. Suspensions where then aliquoted in triplicate over an oil mixture (d = 1.03) prepared from eight parts Silicone Oil (Sigma, Cat # 175633, d = 1.05) and Silicone oil AR20 (Sigma, Cat # 10836, d = 0.95). The cells were then separated from the media and extracted as previously described^40^. Twenty microliters of the extracts and standard aerocavin solutions were analyzed by LC/MS using a Phenomenex Kinetex EVO C18 100Å (100 x 4.6 mm) holding at 65% ACN + 0.1 % FA/water + 0.1% FA with constant flow rate of 0.4 mL/minute.

## ASSOCIATED CONTENT

## Supporting Information

The following files are available free of charge:

Supplementary tables and methods (PDF)

NMR data (PDF)

Accession numbers for analyzed *N. gonorrhoeae* genomes (PDF)

## AUTHOR INFORMATION

## Author Contributions

The manuscript was written through contributions of all authors. All authors have given approval to the final version of the manuscript.

