## Supplementary Tables and Methods for "Aerocavin is an antibiotic with potent and specific anti-Neisserial activity"

Table S1. *S. aureus* mutants resistant to aerocavin

| **Isolate** | **Gene** | **Mutation** |
|---|---|---|
| GP1 | *rpoC* | V1169E |
| GP2 | *rpoB* | L1096F |
| GP3 | *rpoB* | L1135F |
| GP4 | *rpoC* | L1150F |
| GP5 | *rpoC* | I1170T |
| GP6 | *rpoB* | P1125L |
| GP7 | *rpoC* | K334N |
| GP8 | *rpoC* | K334N |
| GP9 | *rpoC* | L1150F |

Table S2. Mutations identified in aerocavin broth passages of *N. gonorrhoeae*

| Line_Passage # | Gene | Mutation | Increase in MIC over the wildtype |
|---|---|---|---|
| 1_28 | *mtrR* | Y136* | 8 |
| | ***rpoB*** | **D1347Y** | |
| | ***rpoB*** | **T1305I** | |
| | *spoT* | D11A | |
| 1_40 | *rpoD* | P555S | 32 |
| | *mtrR* | G/A (promoter) | |
| | *mtrR* | Y136* | |
| | ***rpoB*** | **D1347Y** | |
| | ***rpoB*** | **T1305I** | |
| | *spoT* | D11A | |
| 2_40 | *pilQ* | V115fs | 4 |
| | *spoT* | N252K | |
| | *mtrR* | M1fs | |
| | *atpB* | C89fs | |
| 3_40 | *pilQ* | V162FS | 8 |
| | *spoT* | N252K | |
| | ***rpoB*** | **P1370L** | |
| | *atpA* | P280L | |

Table S3. Strains and growth conditions used in the study

| **Species/strain** | **Propagation** | **MIC testing** | **Conditions** |
|---|---|---|---|
| *Chromobacterium sp.* ATCC 53434 | LB + Amp | CAMHB | 30 °C |
| *Neisseria gonorrhoeae* (all strains) | CHOC | GC | 37 °C, 5% $CO_2$ |
| *Staphylococcus aureus* (all strains) | TS | CAMHB | 37 °C |
| *Escherichia coli* K-12 | LB | CAMHB | 37 °C |
| *Escherichia coli* K-12 *tolC::Kan* | LB + Kan | CAMHB | 37 °C |
| *Acinetobacter baumannii* ATCC 17978 | LB | CAMHB | 37 °C |
| *Pseudomonas aeruginosa* PA14 | LB | CAMHB | 37 °C |
| *Klebsiella pneumoniae* LM21 | LB | CAMHB | 37 °C |
| *Proteus mirabilis* ATCC 29006 | LB | CAMHB | 37 °C |
| *Serratia marcescens* M6C.2 | LB | CAMHB | 37 °C |
| *Salmonella enterica* ATCC 14028 | LB | CAMHB | 37 °C |
| *Mycobacterium smegmatis* mc$^2$ 155 | LB | CAMHB | 37 °C |
| *Enterococcus faecalis* OG1RF | BHI | CAMHB | 37 °C |
| *Staphylococcus epidermidis* ATCC 12228 | BHI | CAMHB | 37 °C |
| *Chromobacterium violaceum* ATCC 12472 | LB + Amp | CAMHB | 30 °C |
| *Shewanella oneidensis* MR-1 | LB | CAMHB | 30 °C |
| *Vibrio cholerae* V52 | LB | CAMHB | 37 °C |
| *Burkholderia cenocepacia* K56-2 | LB | CAMHB | 37 °C |
| *Clostridium perfringens* HN13 | BHIS | BRU | 37 °C, anaerobic |
| *Clostridium difficile* str. 630 | BHIS | BRU | |
| *Neisseria meningitidis* ATCC 13077 | CHOC | MHAB | 37 °C, 5% $CO_2$ |
| *Bacillus subtilis* 168 | LB | CAMHB | 37 °C |

Table S4. Oligonucleotides used in the study

| **Primer** | **Sequence** |
|---|---|
| acvA1_fwd | ccgaagctagcgaattcgtg-AAGGGCATCTCCGAGTGG |
| acvA1_rev | gttcccattt-TCGACTACCTTCCTCAATACAAAG |
| acvA2_fwd | aggtagtcga-AAATGGGAACGCTGCTCTTCC |
| acvA2_rev | aagccggctggcgccaagct-TGCCGCGTCATACTCGATG |
| pNPTS138_rev | CACGAATTCGCTAGCTTCGGC |
| pNPTS138_fwd | AGCTTGGCGCCAGCCGGC |
| pNPTS_sacB_PCR1 | ACAACCATACGCTGAGAGATCC |
| pNPTS_sacB_PCR2 | TTTTACAACGTCGTGACTGG |
| M13_pUC_fwd | CCCAGTCACGACGTTGTAAAACG |
| M13_pUC_rev | AGCGGATAACAATTTCACACAGG |
| acvA-seq1-1 | CACCACCAACAACCGCATGG |
| acvA-seq4 | ACAGCGTCATCGAGCCTATC |

### Supplementary Methods

#### Expression and purification of aerocavin

*Chromobacterium sp.* ATCC 53434 was grown at 30 °C with shaking at 200 rpm in 5 mL LB broth overnight. The cultures were then spread on the following medium (adapted from Singh *et al*[1] ): 10 g/L yeast extract, 10 g/L malt extract, 1 g/L peptone, 20 g/L glucose, 15 g/L agar. The cultures were incubated at room temperature for five days, and then the agar was extracted three times with ethyl acetate. Ethyl acetate was removed by evaporation under vacuum, and the crude extract was solubilized in chloroform and extracted three times with water. Chloroform was removed by evaporation, and the extract was fractionated with C18 Sep-Pak columns (Waters) with stepwise acetonitrile gradient. The active fractions were further separated with Silica Sep-Pak columns (Waters) using stepwise hexanes/ethyl acetate gradient. The fraction containing the activity was injected into Phenomenex Synergi 4 µm Hydro-RP 80 Å, 250 x 10 mm preparative column holding 65% acetonitrile + 0.1% formate at 10 mL/minute flow rate. Purified aerocavin was stored at -20 °C in methanol.

#### NMR spectroscopy

The product's structure was confirmed with NMR. NMR spectral data were obtained by using a Bruker Avance 600 MHz spectrometer (Bruker), and the chemical shift values were represented in δ scales. Residual protium and carbon resonances of $CDCl_3$ NMR solvent were used to reference chemical shifts ($\delta_H$ 7.25 and $\delta_C$ 77.0). All the one-bond $^1H$–$^{13}C$ correlations were identified based on $^1H$, $^{13}C$, and HSQC NMR data, and the planar structure was assigned by detailed analysis of

COSY NMR spectrum along with interpretation of two- and three-bond $^{1}H-^{13}C$ HMBC correlations.

**Paper disk activity screen**

*N. gonorrhoeae* was grown overnight on chocolate agar, scraped, resuspended in Mueller Hinton broth, and spread with cotton-tipped applicators onto GCB agar supplemented with + 1% IsoVitaleX™ solution (BD). All other species were grown overnight in Mueller-Hinton broth and spread onto Mueller-Hinton Agar. Paper disks (Thermo) were impregnated with 100 μg of aerocavin and placed on top of the agar. *N. gonorrhoeae* was incubated at 37 °C in 5% $CO_2$, the other species were incubated under atmospheric conditions at 37 °C. *M smegmatis* was incubated for 48 hours, the other the species were incubated for 24 hours. To control for differences that media composition and atmospheric $CO_2$ concentration may have on aerocavin's activity, *S. aureus* Newman was additionally tested under the conditions used for *N. gonorrhea* to ensure that the results were comparable.

**Determination of minimal inhibitory concentrations**

Minimal inhibitory concentrations (MICs) were measured following CLSI guidelines[2-4]. MICs of *Neisseria* and *Clostridia* were determined using agar dilution assays[3, 4]. MICs of other bacteria were determined using broth macrodilution assays with agitation at 200 r. p. m. Table S3 contains information on media and conditions used for each strain. In all cases *S. aureus* Newman was included to control for large effects of differing conditions on aerocavin's activity. Sixty-four μg/mL was the highest concentration tested due to limited solubility of aerocavin.

**Presence of aerocavin resistance-associated mutations in clinical *N. gonorrhoeae* isolates.**

Publicly-available whole genome sequencing reads from 12,086 N. gonorrhoeae clinical isolates were downloaded from the European Nucleotide Archive[5-27]. Reads were assembled with SPAdes v 3.12.0[28] using the careful flag to correct assembled contigs. We additionally removed any contigs less than 500 nucleotides in length or with less than 10x coverage. We used blastn from BLAST+ v 2.9.0[29] to identify sequences of genes encoding the RNA polymerase in assembled genomes. Translated amino acid sequences were aligned with MAFFT v 7.402[30]. We characterized the diversity at each amino acid position previously identified to be associated with aerocavin resistance in other organisms.

**Isolation of resistant mutants**

*S. aureus*. Individual colonies were inoculated into tryptic soy broth (TSB) and incubated overnight at 37 °C with shaking at 200 rotations per minute (rpm). These cultures were then diluted 1/20 into liquid TSB containing 4X MIC of aerocavin and incubated overnight producing growth. The cultures were struck out on antibiotic-free media for acquisition of individual colonies and confirmed to retain aerocavin resistance after passages in antibiotic free-media.

*N. gonorrhoeae.* Colonies of resistant mutants were picked from the plates that were used to determine the frequency of resistance. The resistant mutants were re-streaked on agar containing aerocavin at the same concentration as that at which they were selected. To acquire isolates with secondary mutations, the procedure was repeated to isolate step 2 and step 3 mutants.

We did not isolate mutants from liquid passages of *N. gonorrhoeae*. Instead, passages 28 and 40 from line 1, and passages 40 from line 2 and 3 were spread from the frozen stock onto chocolate agar plates and incubated overnight. The resulting biomass was collected for DNA isolation and sequencing.

**DNA sequencing**

DNA from resistant mutants was purified using Wizard® Genomic DNA Purification Kit (Promerga) or ZymoBIOMICS DNA Microprep Kit (Zymo Research) followed by cleanup with Genomic DNA Clean & Copncentrator-10 (Zymo Research). Illumina libraries were prepared using NEBNext® Ultra$^{TM}$ DNA Library Kit (New England Biolabs) and sequenced on a MiSeq System with MiSeq Reagent v3 (2 x 300 bp) Kit (Illumina, Inc.). The data were analyzed using CLC Genomics software (QIAGEN).

**RNA polymerase inhibition assay**

RNA polymerase inhibition assay was carried out using *E. coli* RNA Polymerase Kit (Profoldin) and *E. coli* RNA Polymerase Holoenzyme (NEB). The assays were carried out in 384-well black flat bottom plates (Corning) adhering to the RNA Polymerase Kit instructions except RNAP purchased from NEB was used instead of the RNAP supplied with the kit. Prior to addition of the DNA template, the reactions were incubated with aerocavin or vehicle (methanol) for 5 minutes at 37 °C. The fluorescent readout of RNAP activity was measured on CLARIOstar multimode plate reader (BMG Biotech) with 485 nm excitation and 535 nm emission wavelengths. The data were analyzed using GraphPad Prism 10 using non-linear fit.

(2) CLSI. *Performance Standards for Antimicrobial Susceptibility Testing. 31st ed. CLSI Supplement M100. Clinical and Laboratory Standards Institute*; 2021.

(3) CLSI. *Methods for Antimicrobial Susceptibility Testing of Anaerobic Bacteria. 9th ed. CLSI standard M11.* ; Wayne, PA, 2018.

(4) CLSI. *Methods for Dilution Antimicrobial Susceptibility Tests for Bacteria That Grow Aerobically. 11th ed. CLSI Standard M07*; Wayne, PA, 2018.

(18) Lan, P. T.; Golparian, D.; Ringlander, J.; Van Hung, L.; Van Thuong, N.; Unemo, M. Genomic analysis and antimicrobial resistance of Neisseria gonorrhoeae isolates from Vietnam in

2011 and 2015-16. *J Antimicrob Chemother* **2020**, *75* (6), 1432-1438. DOI: 10.1093/jac/dkaa040 From NLM Medline.
